# Velocity estimation in reinforcement learning

**DOI:** 10.1101/432492

**Authors:** Carlos A. Velázquez, Manuel Villareal, Arturo Bouzas

## Abstract

The current work aims to study how people make predictions, under a reinforcement learning framework, in an environment that fluctuates from trial to trial and is corrupted with Gaussian noise. A computer-based experiment was developed where subjects were required to predict the future location of a spaceship that orbited around planet Earth. Its position was sampled from a Gaussian distribution with the mean changing at a variable velocity and four different values of variance that defined our signal-to-noise conditions. Three error-driven algorithms using a Bayesian approach were proposed as candidates to describe our data. The first is the standard delta-rule. The second and third models are delta rules incorporating a velocity component which is updated using prediction errors. The third model additionally assumes a hierarchical structure where individual learning rates for velocity and decision noise come from Gaussian distributions with means following a hyperbolic function. We used leave-one-out cross-validation and the Widely Applicable Information Criterion to compare the predictive accuracy of these models. In general, our results provided evidence in favor of the hierarchical model and highlight two main conclusions. First, when facing an environment that fluctuates from trial to trial, people can learn to estimate its velocity to make predictions. Second, learning rates for velocity and decision noise are influenced by uncertainty constraints represented by the signal-to-noise ratio. This higher order control was modeled using a hierarchical structure, which qualitatively accounts for individual variability and is able to generalize and make predictions about new subjects on each experimental condition.

## Introduction

Decisions often take place in environments that change over time. Availability of food in foraging animals may vary gradually as a function of source growth or continuous intake, likewise, the position of objects in space can change at a certain rate. Having accurate estimates of relevant variables under these circumstances allows behavior to be better allocated. For example, by changing to a richer foraging location or predicting the correct position of an object moving towards us. In stable environments, prediction error models accomplish this task by reducing the discrepancy of estimates and outcomes as new observations arrive. A straightforward expression to compute this is the Delta-rule:

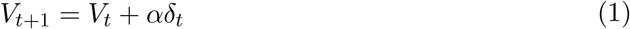

where, the estimate at time *t* + 1 (*V*_*t*+1_), depends on the previous estimate (*V_t_*) and the prediction error (*δ_t_*), weighted by the learning rate parameter (*α*). Evidence from Experimental Psychology (Bush & Mosteller, 1951; Dayan & Nakahara, 2018; Miller, Barnet, & Grahame, 1995; Rescorla, Wagner, et al., 1972) and Neuroscience (Daw & Tobler, 2014; Niv, 2009; Schultz, Dayan, & Montague, 1997) provides support for this algorithm as a plausible mechanism of learning in mammals, and it has also been implemented as an effective solution in multiple machine learning problems (Sutton & Barto, 1998). However, one of its limitations is the inability to describe behavior in non-stationary environments, partly, due to the fixed nature of the learning rate parameter (O’Reilly, 2013). For example, in change-point problems, having a low *α* makes predictions during stable periods accurate but causes a slow adaptation after a change. A high α has the opposite effect, making inaccurate predictions during stability but having a quick adaptation to changes. Adjusting this parameter after the change-point (Nassar, Wilson, Heasly, & Gold, 2010) and using multiple delta-rules with their own learning rates (Wilson, Nassar, & Gold, 2013) are some of the possible solutions that have been proposed. On the other hand, when the environment changes gradually over trials such as in a random walk process, the learning rate is assumed to vary as function of the relative uncertainty in the estimates and the outcomes as expressed in the Kalman filter equations (Gershman, 2015, 2017; Kakade & Dayan, 2002; Kalman, 1960; Navarro, Tran, & Baz, 2018; Speekenbrink & Konstantinidis, 2015; Speekenbrink & Shanks, 2010; Zajkowski, Kossut, & Wilson, 2017). An important limitation of this approach is that, when trial-to-trial changes are large (i.e. the rate of change is high), the learning rate asymptotes at values close to one (Daw & Tobler, 2014), making the model extremely sensitive to outcome noise. This problem is likely to occur because there is not an explicit computation of the rate of change of the environment.

In this work we show that when the environment is changing at certain rate, a concrete estimation of this variable is necessary to guide decisions. Additionally, we show that the updating process of the rate of change is influenced by the level of noise in the observations as expressed by the signal-to-noise ratio (S/N). Previous research has shown that people are sensitive to higher order statistics of the environment such as the volatility (Behrens, Woolrich, Walton, & Rushworth, 2007; O’Reilly, 2013) or the functions controlling changes (Ricci & Gallistel, 2017) and that they are able to adapt their behavior accordingly.

In our experiment, subjects were required to predict the angular location of a spaceship moving around planet Earth. Its position was generated from a Gaussian distribution with the mean changing at a variable velocity (rate of change for position) and four values of variance that defined the S/N conditions. We proposed a reinforcement learning model incorporating a velocity component to describe participants’ predictions throughout the task. The main assumption of the model is that prediction errors are used to update an estimate of the velocity in the generative process, which is later incorporated to the computations of new predictions.

A hierarchical extension of the model was implemented where individual parameters were generated from Gaussian distributions defined at the level of conditions. In general, hierarchical modeling allows to specify the generative process of relevant psychological variables rather than assuming they simply exist (Lee, 2018; Shiffrin, Lee, Kim, & Wagenmakers, 2008). One of the main advantages of this type of models is their ability to generalize the results to new conditions or participants (Lee, 2018). In the current work, we assumed that the means of the Gaussian distributions for two of the model parameters (the rate of learning for the velocity component and the decision noise) followed a hyperbolic function of the S/N values. These assumptions suggest that the overall behavior of these parameters is sensitive to the level of noise in the observations.

Our results show that errors between the generative mean of the spaceship and participants’ predictions remain close to zero in the four conditions and that accuracy increases with the S/N. The model-based analysis indicates that a prediction error model incorporating a velocity component is better at describing participants’ behavior in the task than model that ignores the computation of this variable. Additionally, we found that a hierarchical extension is able to make sensible predictions about a new participant in the four conditions and potentially to new S/N values. A formal model comparison, using a new approach to leave-one-out cross validation developed by Vehtari, Gelman, and Gabry (2017) and the Widely Applicable Information Criterion (WAIC), suggests that the hierarchical model has the best predictive power. We further discuss the implications of our findings for reinforcement learning models and alternative approaches to similar prediction problems.

## Learning models

We evaluated three error-driven algorithms using a graphical Bayesian approach (Farrell & Lewandowsky, 2018; Lee & Wagenmakers, 2013). The first, is the standard delta-rule of equation 1 (Bush & Mosteller, 1951; Dayan & Nakahara, 2018; Miller et al., 1995; Rescorla et al., 1972). The second and third models, correspond to a delta-rule that incorporates a velocity term. The key difference between them is that the later assumes a hierarchical structure (Lee, 2018; Shiffrin et al., 2008) where parameters come from Gaussian distributions.

### Standard delta-rule (SD)

Equation 1 specifies the structure of this model. We assumed a decision rule where responses of participants for the next trial, *B*_*t*+1_, were generated from a Gaussian distribution with mean *V*_*t*+1_ and variance *η* representing decision noise, i.e.,

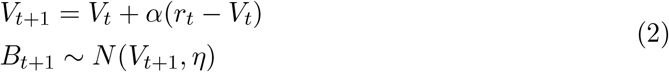

Figure 1 is the graphical representation of this model. In this notation, nodes correspond to variables and arrows connecting them refer to dependencies. Shaded nodes are observed variables, whereas unshaded nodes are latent variables. Stochastic and deterministic variables are represented using single- and double-boarded nodes, respectively, and continuous variables are represented using circular nodes. Plates refer to replications of the process inside them. On the right-hand side of the graphic, we show the detailed relations among variables and the prior distributions of the parameters. The learning rate *α* is assumed to be within the interval (0,1) as it is commonly specified for prediction error models. A uniform distribution over this range indicates that there is no specific value of the parameter that is more likely before data is observed; the parameter *η* is assumed to have the same prior distribution as *α*. Finally, note that we used precision (inverse of *η*) to specify the error term of the Gaussian distribution from which responses are generated. This modification was done only due to code specifications for the analysis.

**Figure 1.**
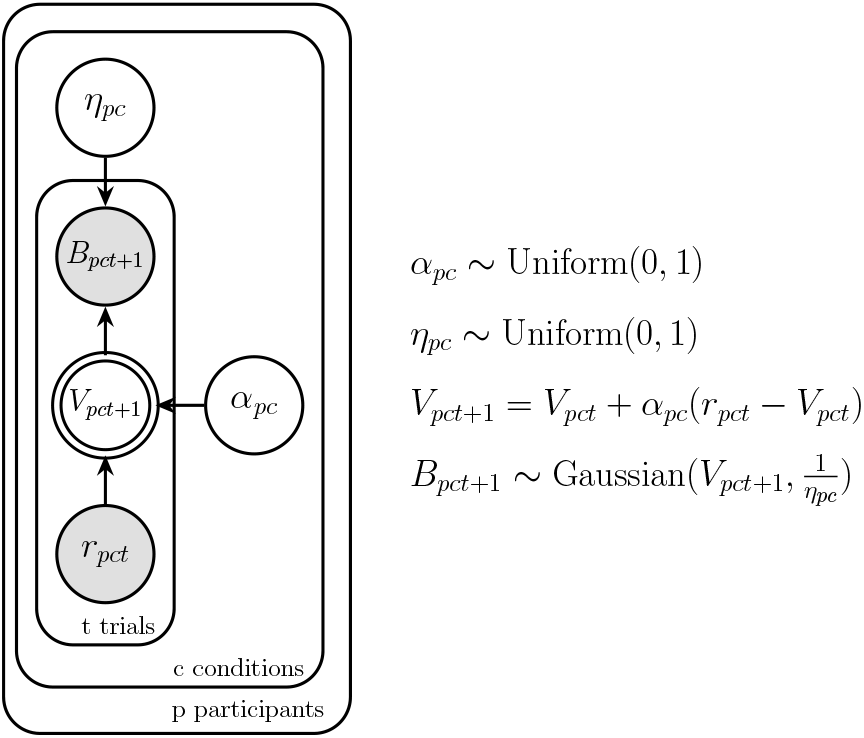
Graphical representation of delta-rule.

Although a very useful model of animal and machine learning, the core structure of equation 3 has difficulties performing under changing conditions (Gallistel, Krishan, Liu, Miller, & Latham, 2014; Nassar et al., 2010; Ricci & Gallistel, 2017; Ritz, Nassar, Frank, & Shenhav, 2018; Wilson et al., 2013). In particular, for the purpose of this work, we emphasize that it does not compute a potential rate of change in the environment.

### Delta-rule with velocity term (VD)

This model is derived from the formal structure of a Kalman filter incorporating a velocity term (Kalman, 1960; Ritz et al., 2018). The Kalman filter represents an idealized Bayesian learner that updates estimates about a variable of interest that is changing over trials and is corrupted with Gaussian noise. We simplified the model by assuming that learning rates (known as the “gain”) and decision noise are free parameters for each condition. Our analysis was based on this assumption given that in non-stationary environments like ours, learning rates stabilize in values that asymptotically correspond to the free parameters (Daw & Tobler, 2014). Furthermore, our analysis focused on the overall behavior of parameters rather than on the dynamics as specified in Kalman filter equations. Formally, this model can be expressed as delta-rule with a velocity term:

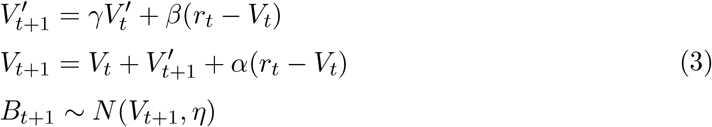

where 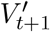 is the predicted velocity for trial *t* +1 and is updated according to its own delta-rule with learning rate *β*. *γ* is a memory retention parameter that indicates the weight of the previous velocity in the updating process. Note that if 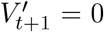 the model reduces to equation 2. Figure 2 shows the graphical representation of the VD model. In the same way as *α* in the previous model, *β* and *γ* are assumed to be uniformly distributed within the interval (0,1) before observing the data. Apart from that, specifications of the graphical model in figure 2 are the same as in figure 1.

**Figure 2.**
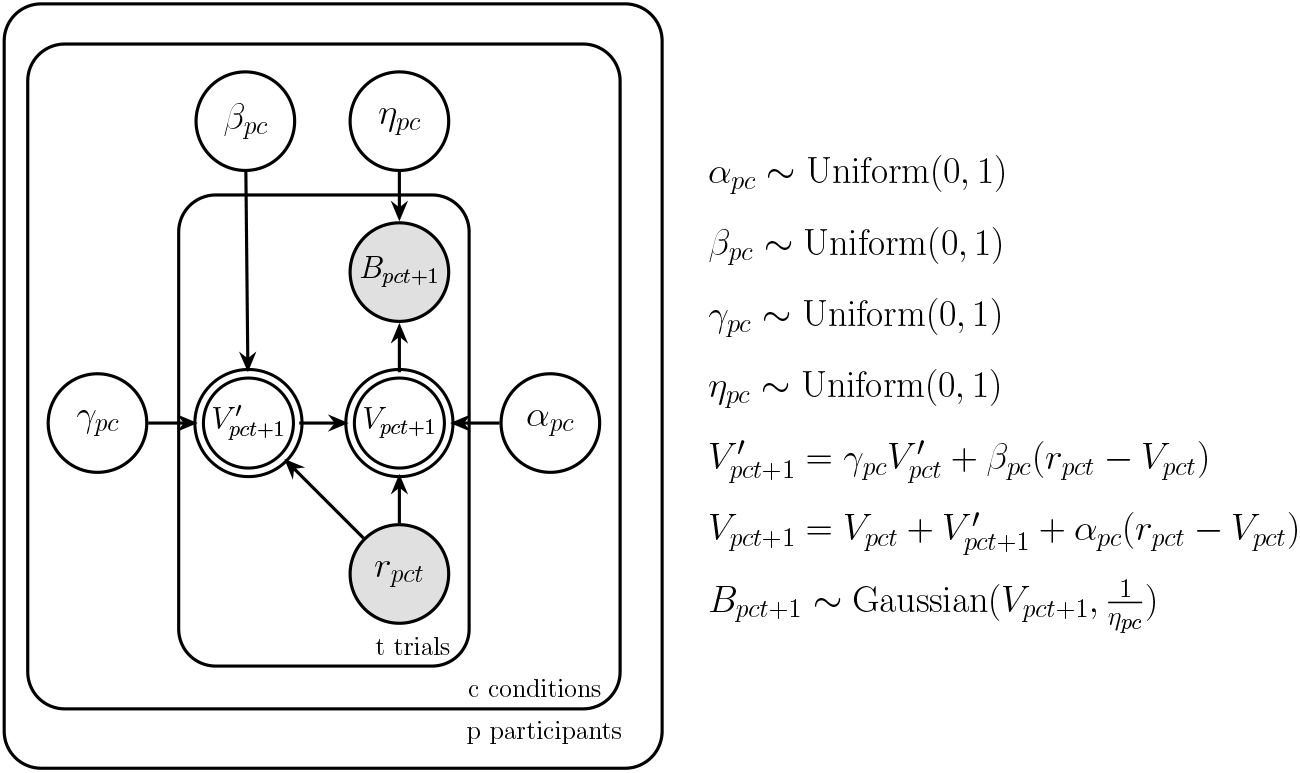
Graphical representation of delta-rule with a velocity term.

### Hierarchical delta-rule with velocity term (HVD)

This model is a hierarchical extension of VD. Hierarchical models assume that individual parameters are themselves the outcome of a higher-order process (Lee, 2018; Shiffrin et al., 2008). Some of their applications involve modeling individual differences assuming participants are not completely independent (Pratte & Rouder, 2011), and predicting behavior of a new participant based on information of the population (Lee, 2018; Shiffrin et al., 2008). One simple way to express a hierarchy, is to set Gaussian distributions from which parameters are generated (Matzke, Dolan, Batchelder, & Wagenmakers, 2015; Zajkowski et al., 2017). Figure 3 is a graphical representation of this model. Note that individual parameters now depend on a mean *μ* and precision 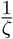 (where *ζ* is the variance of the distribution) commonly known as hyperparameters. The above implies that what is learned from one participant in a given condition affects what is learned about the rest in the same condition. In particular, after observing the results of model VD, we decided that values for the learning rates for position α and the memory retention parameter *γ* could be generated by Gaussian distributions for the whole experiment. This assumption is based on the fact that these parameters barely changed from one condition to the other. However, in the case of learning rates for velocity *β* and decision noise *η*, we assumed they could be generated from Gaussian distributions with means following a hyperbolic function. As will be described below, this assumption arises from visual inspection of individual parameters and the behavior of the model when tracking the generative process of the experiment. Each hyperbola consists of the S/N value as argument and two parameters with positive values controlling the shape of the function (*a^β^* and *b^β^* for learning rates of velocity and *a^η^* and *b^η^* for decision noise). We specified uniform prior distributions over the range (0,1) for the majority of hyperparameters, with the exception of *b^β^, a^η^* and *b^η^* as they can take values higher than one. For these hyperparameters we set non-informative priors represented by truncated Gaussian distributions with a low precision.

**Figure 3.**
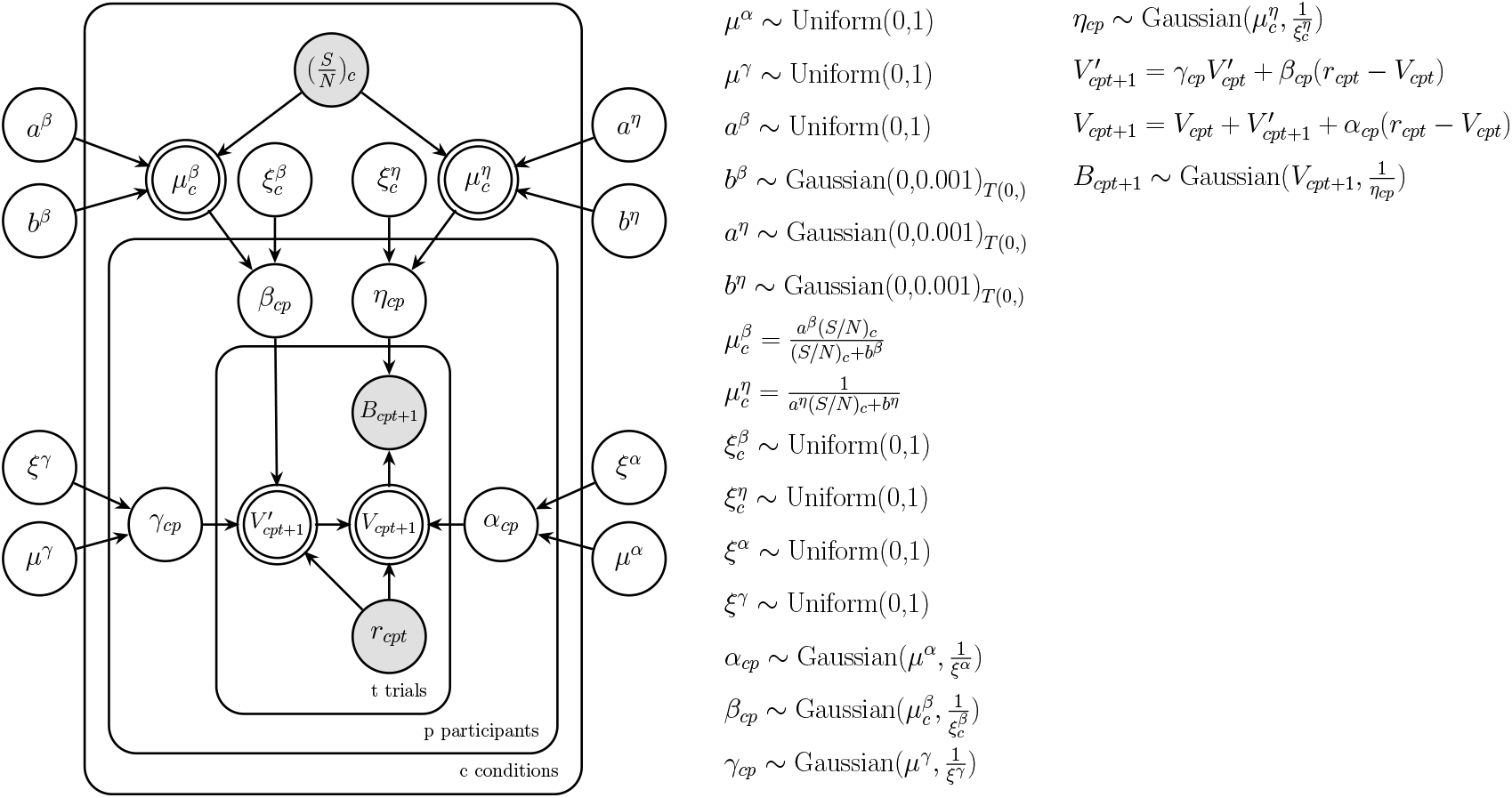
Graphical representation of a hierarchical delta-rule with a velocity term.

## Method

### Participants

Seventy two undergraduate students (55 female, mean (SD) age = 19.8 (2.03)) from the School of Psychology at the National Autonomous University of Mexico participated in the study after providing informed consent.

### Behavioral task

The experiment was programmed using the commercial software Matlab and the extension Psychophysics Toolbox (Brainard & Vision, 1997; Kleiner et al., 2007; Pelli, 1997) to create visual stimuli. A standard mouse and keyboard, and a screen of 1920 × 1080 pixels were used.

Figure 4 displays a representation of the task. In this scenario, participants were required to predict the future position of a spaceship moving around planet Earth along a specified orbit (blue dotted circle). Its location was given in radians (rad) using the center of the orbit as reference. On every trial, the spaceship appeared for half a second on a given point of its orbit, after which it disappeared. Participants had to click on the position where they thought it would reappear in the next trial. After their choice, a red dot indicated the location they selected and a red circle a fixed margin of error. At the same time, the spaceship showed up in its new position. If it fell inside the red circle, it turned red indicating the new location was accurately predicted or remained the same color otherwise. There was no time limit for participants to emit their responses. This sequence was repeated throughout the experiment. Moreover, if the spaceship gave a full lap in a counterclockwise direction, i.e. moving 2*π* rad, the second lap would continue from 2*π* rad to 4*π* rad, and so on. Similarly, if the spaceship completed a full lap in a clockwise direction, its next position was given according to values of the previous lap. We followed the same logic to register participant’s responses. This transformation allowed the range of possible values of observations and responses to span from −∞ to ∞, and was particularly useful to avoid sudden changes of position from 2*π* rad to 0 every time a full lap was completed.

**Figure 4.**
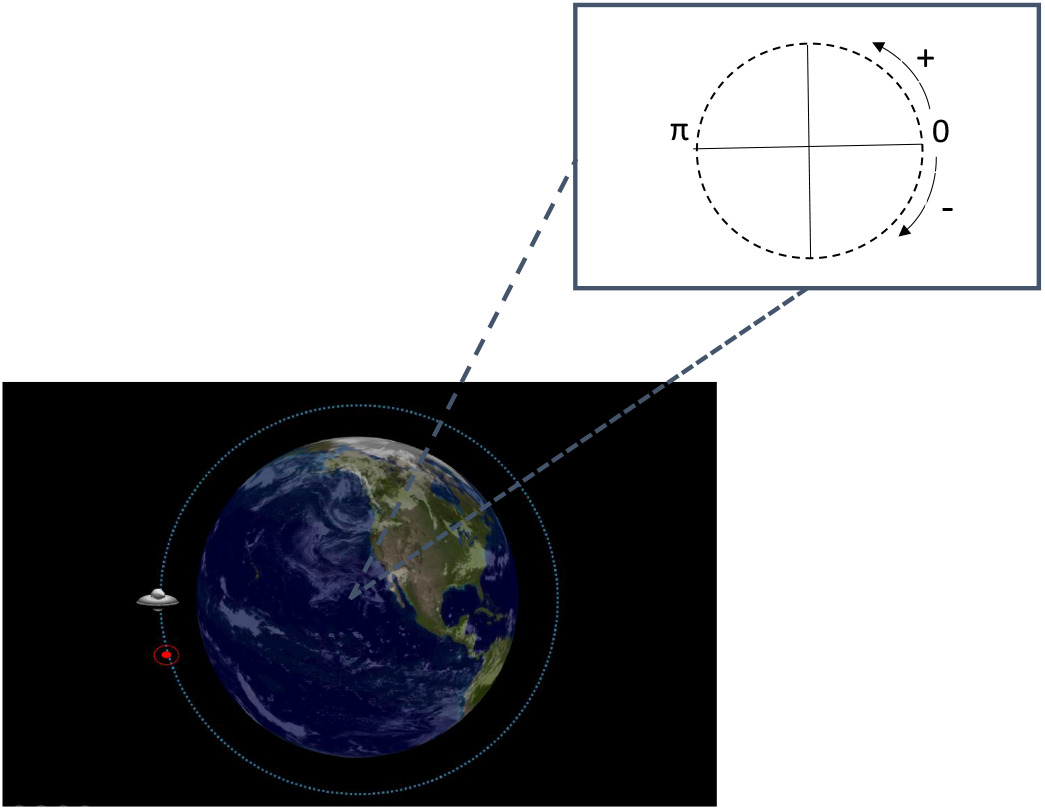
Representation of the behavioral task. Participants predicted the position of a spaceship that moved around planet Earth along the blue dotted line. Selected positions were indicated with a red circle surrounded by a red circle representing a margin of error. Graphics of the task were developed using the on-line site http://planetmaker.wthr.us/. See the main text for detailed explanation of the task.

### Experimental design

On every trial *t* the position of the spaceship *r* was generated from a moving Gaussian distribution following:

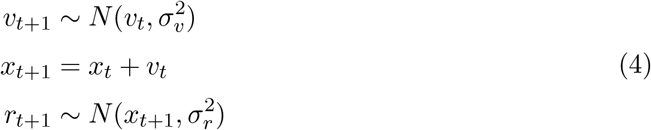

where *x* is the mean of the distribution, *v* is a velocity term following a Gaussian random walk with variance 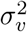, and 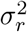 is the variance in the actual observations. Our experiment consisted of four conditions that varied the S/N values represented by 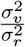. Intuitively, this quantity indicates how easy is to discriminate changes due to velocity, relative to changes due to random noise; smaller ratios indicate noisier observations. We fixed the numerator of the ratio 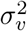 so participants faced the same generative process for the velocity component, but varied the denominator 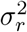 to change the noise in their observations. Table 1 shows the values of S/N used in the experiment. An experimental session consisted of four conditions with 300 trials each and order of presentation was randomized for all participants. Before the experimental task started, a practice phase was completed that consisted of at least 30 trials, after which participants decided whether to continue acquiring more practice or begin the experiment.

**Table 1.** Experimental conditions. Units of S and N are given in rad^2^

| Condition | $\frac{S}{N}$ | $S$ | $N$ | Trials |
| --- | --- | --- | --- | --- |
| 1 | 0.05 | 0.0049 | 0.098 | 300 |
| 2 | 0.5 | 0.0049 | 0.0098 | 300 |
| 3 | 1 | 0.0049 | 0.0049 | 300 |
| 4 | 2 | 0.0049 | 0.00245 | 300 |

## Results

As a measurement of performance we report the error between the generative mean of the spaceship and participants’ predictions. These values are shown in the bottom panels of figure 5 for all subjects on the four S/N conditions. It is clear that, in all conditions, most errors remain close to zero, however, accuracy increases with the S/N. This is evident when observing the proportion of values around zero for each condition and the corresponding standard deviation. In other words, participants had greater errors for noisier observations. Top panels are graphical representations of bottom plots where dark and light blue represent ±*σ* and ±2*σ*, respectively, and the figure of the spaceship represents the generative mean. For individual errors in each condition see supplementary material (see Supplementary material).

**Figure 5.**
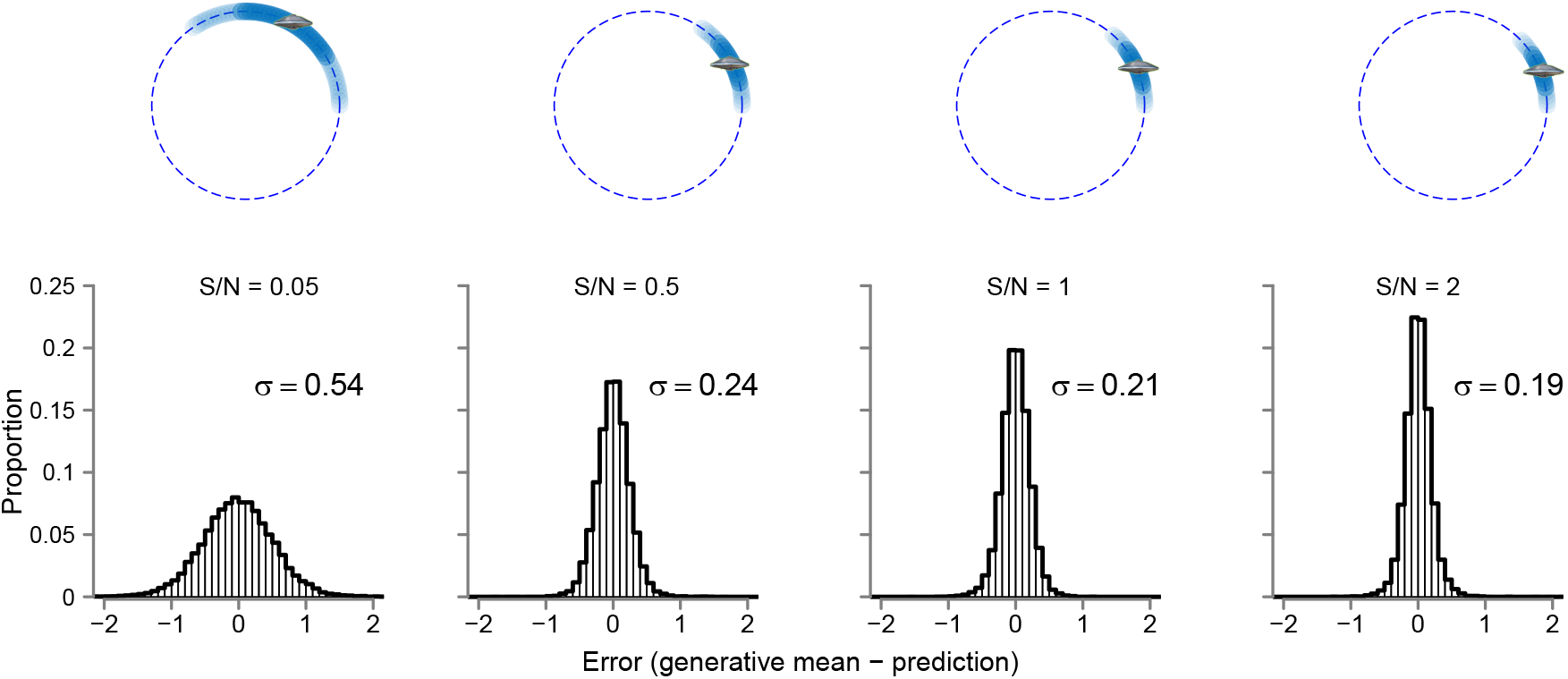
Overall performance of subjects in the experiment. Bottom panels show the error between the generative mean of the spaceship and the predictions of participants for all trials on each of the conditions indicated with their corresponding S/N values. A graphical representation of the errors is shown in the top panels where dark and light blue indicate ±*σ* and ±2*σ*, respectively, and the spaceship represents the generative mean.

## Bayesian Inference

Posterior distributions of parameters and hyperparameters were approximated using the software JAGS (Just Another Gibbs Sampler; Plummer 2003) implemented in R code. This procedure uses a sampling method known as Markov chain Monte Carlo (MCMC) to estimate parameters in a model. For our three graphical models we used three independent chains with 10^5^ samples each and a burn-in period (samples that were discarded in order for the algorithm to adapt) of 8 × 10^4^. A thinning of 20 was used (i.e., values were taken every 20 samples of the chain) to reduce autocorrelation within chains. Convergence was verified by computing the *Ȓ* statistic (Gelman, Rubin, et al., 1992) which is a measure of between-chain to within-chain variance where values close to 1 indicate convergence (Lee & Wagenmakers, 2013). In general, values between 1 and 1.05 are considered as reliable evidence for convergence. All of the nodes in SD and HVD models and around 99% of the nodes in VD model had *Ȓ* values within this interval.

Figure 6 shows the results for the SD (gray) and VD (blue) models. The four left panels display maximum posterior values (modes) of parameters for each participant, ordered by experimental condition (S/N values). Error bars correspond to the interquartile range and dots represent the medians. It can be observed that values of learning rates for position *α* and decision noise parameters *η* for SD model were higher in all conditions compared to the ones of the VD model. In particular, the values of *α* for the SD model are not visually different from one. The above results suggest that when making predictions participants ignored the history of previous observations and were purely guided by the just-observed outcome. This would imply that their predictions were highly dependent on the level of noise in the condition. Maximum posteriors for the VD model of both *α* and *η* are lower than in SD model in all conditions as a result of incorporating the velocity component. On the other hand, values for the memory retention parameter *γ* are close to one in all conditions indicating that, when updating the velocity term, the estimate of the previous velocity was retained almost perfectly. In the case of the learning rates for velocity *β*, we observe a gradual increment of values as the S/N increases, which suggests that the velocity term was updated faster for less noisy observations. Additionally, it is shown that decision noise *η* for the VD model decreases for higher values of S/N. This relation indicates that when observations were less noisy so were the predictions of participants. The right panel of figure 6 shows the root mean squared error (RMSE) generated from the posterior predictions of each model compared to the actual RMSE of participants. Posterior predictions were obtained by simulating data with 300 samples with replacement from the joint posterior distribution of parameters and the actual observations of participants. The similarity between the RMSE of the simulated data and the actual RMSE is an indicator of the descriptive adequacy of the models. Note that in all conditions the model incorporating the velocity component recovers the actual RMSE better than the SD model.

**Figure 6.**
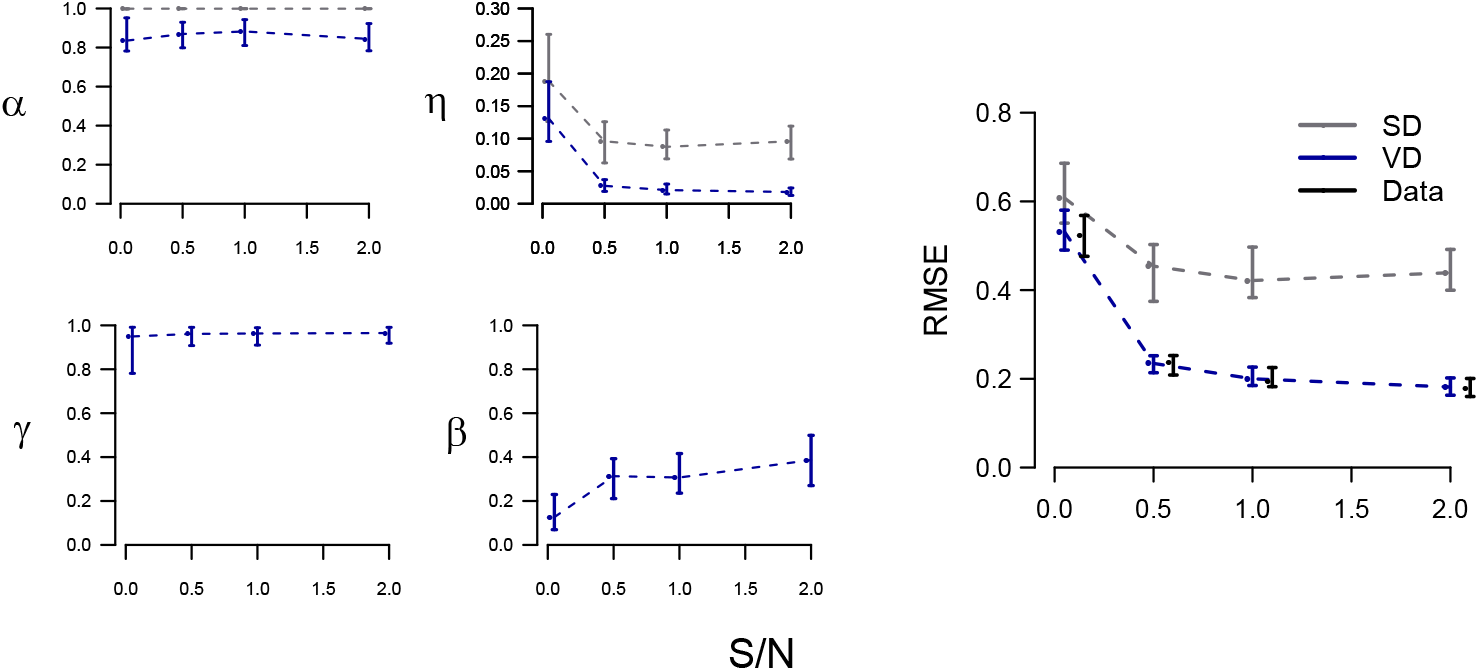
Bayesian inference results for SD and VD models. The four left panels show maximum posteriors for the learning rates of position (*α*), learning rates of velocity (*β*), velocity decay (*γ*) and decision noise (*η*) parameters for all subjects on each condition represented by the corresponding S/N value. Error bars represent the interquartile range and dots the medians. Values for the SD model are shown in gray and those for the VD model in blue. The right panel displays the RMSE generated with the posterior predictions of each model compared to the actual RMSE of participants.

Although VD model qualitatively describes our data better than SD model, neither of them provides information for new data sets or makes predictions for different values of the S/N. These limitations are overcome by assuming a hierarchical structure over the parameter space. By visual inspection of figure 6, we can tell that *α* and *γ* are invariant over different values of the S/N. Thus, we can simplify the model by assuming they are generated by two independent Gaussian distributions for the whole experiment. However, this is not the case for *β* and *η*. These parameters appear to gradually increase and decrease, respectively, as the S/N increases. To formalize this pattern, we assumed that *β* and *η* are generated from Gaussian distributions with means following a hyperbolic function of the S/N values. The graphical model of figure 3 (HVD) specifies each hyperbola. It is important to emphasize that, in addition to visualization of parameters in figure 6, the decision of using of a hyperbolic function was based on the behavior of the parameters when the model was allowed to track the generation process of the experiment (see Supplementary materials). Figure 7 shows the results for the HVD model. Left and middle columns correspond to the means of the Gaussian distributions and represent the overall behavior of individual parameters. In the left column, we show posterior samples of the mean of learning rates for position *μ^α^* (top) and memory retention parameters *μ^γ^* (bottom). As previously stated, they comprise single distributions for the whole experiment and therefore the S/N values were omitted. The middle column corresponds to hyperbolas for the hierarchical means of learning rates for velocity *μ^β^* (top) and decision noise *μ^η^* (bottom). These functions were generated by drawing 300 samples with replacement from the joint posterior distribution of the parameters that constitute each hyperbola (i.e. *a^β^, b^β^* and *a^η^, b^η^*) and values of S/N in steps of 0.01 within the interval (0,2). Importantly, given that the functions are continuous, we are able to make predictions about the average behavior of parameters for untested S/N values. In the right panel of figure 7, we show the descriptive adequacy of the HVD model using the same sampling procedure as for the SD and VD models. It can be observed that model HVD is able to recover the actual RMSE of subjects just as accurately as its non-hierarchical version in figure 6. The ability to capture individual behavior is an important property of hierarchical models (Pratte & Rouder, 2011) and is represented by the variances of the higher-level distributions (see Supplementary materials).

**Figure 7.**
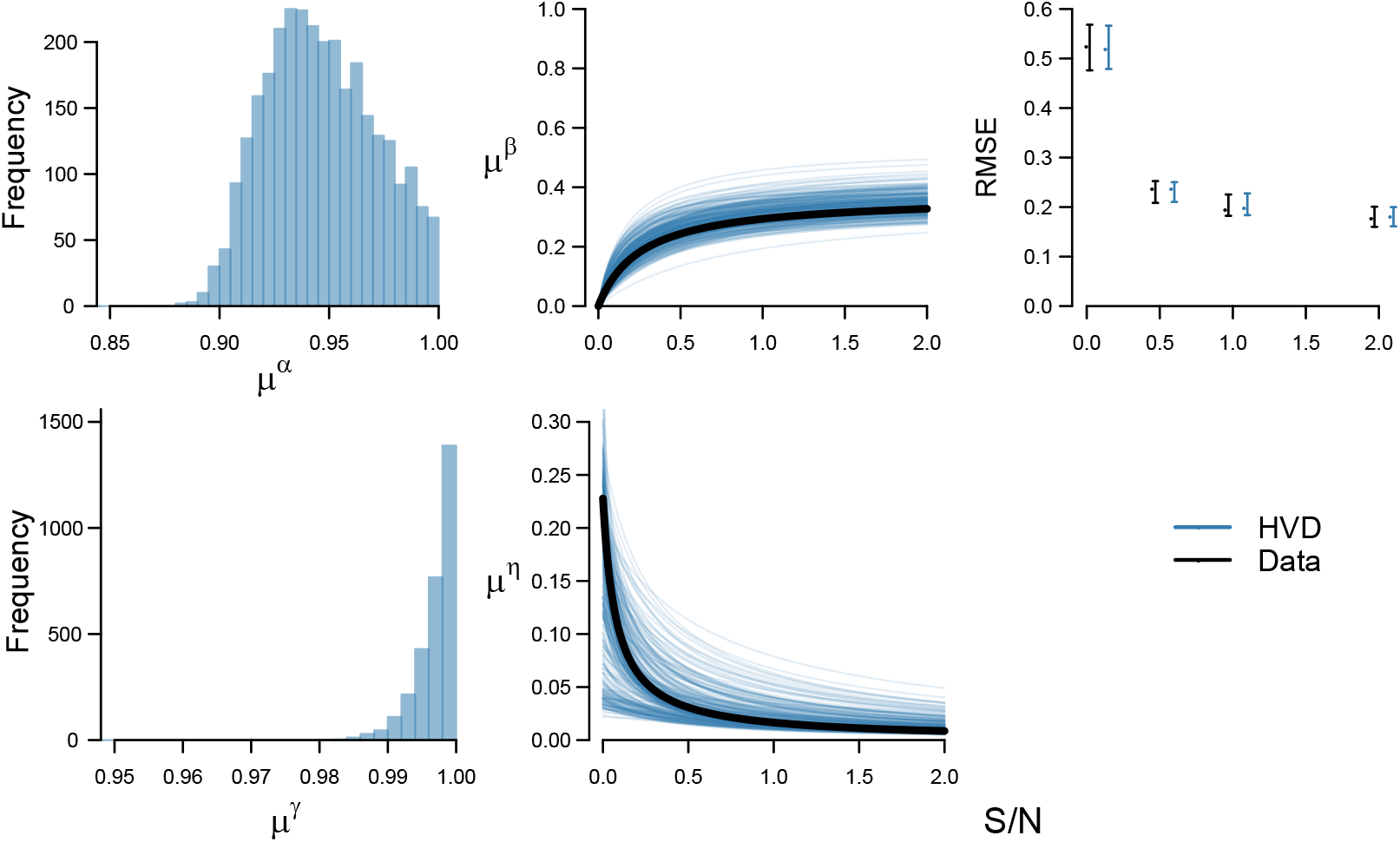
Results of Bayesian inference for the HVD model. The left panels show posterior samples of the hierarchical means for learning rates of position (top) and the memory retention of velocity (bottom). Middle panels correspond to the hyperbolas for the hierarchical means of learning rates of velocity (top) and decision noise parameters (bottom) as a function of the S/N. Each hyperbola was generated by sampling with replacement from the joint posterior distribution of *a^β^* and *b^β^*, for learning rates of velocity means, and *a^η^* and *b^η^*, for decision noise. The black solid line represents the hyperbola generated with the maximum posterior values. The right panel shows the RMSE generated with the posterior predictions of the model for all subjects compared to the actual RMSE of participants. Error bars represent the interquartile range.

An additional benefit from the hierarchical structure of model HVD is shown in figure 8. In this graph, we simulated data for a new participant on each experimental condition. For SD and VD models, predictions were generated from the prior distributions of parameters. This is because in non-hierarchical models participants are assumed to be independent of each other and, therefore, the collected data does not update the beliefs about a new subject. For HVD model, inferences about the parameters of a new participant come from the group-level distributions for each condition, which, in turn, are inferred from data of all participants. Top panels of figure 8 correspond to the predicted RMSEs for the VD and HVD model. Values for the SD model were excluded as they considerably increased the scale of the plot making difficult the visualization of RMSEs of the other two models. It is evident that, although VD accurately described the performance of subjects in the task, it makes very poor predictions about a new participant. Conversely, the HVD model predicts that a new subject would have RMSEs similar to the ones actually observed from participants. Bottom panels in figure 8 show the trial-by-trial predictions of the three models on each experimental condition. The colored fringes correspond to 300 simulations for a new set of observations following the same generative process of the experiment. As can be observed, SD and VD make very broad predictions on every trial compared to the HVD model. However, it is worth noting that simply incorporating the velocity component into the VD model considerably reduces the variability of the predictions of SD.

**Figure 8.**
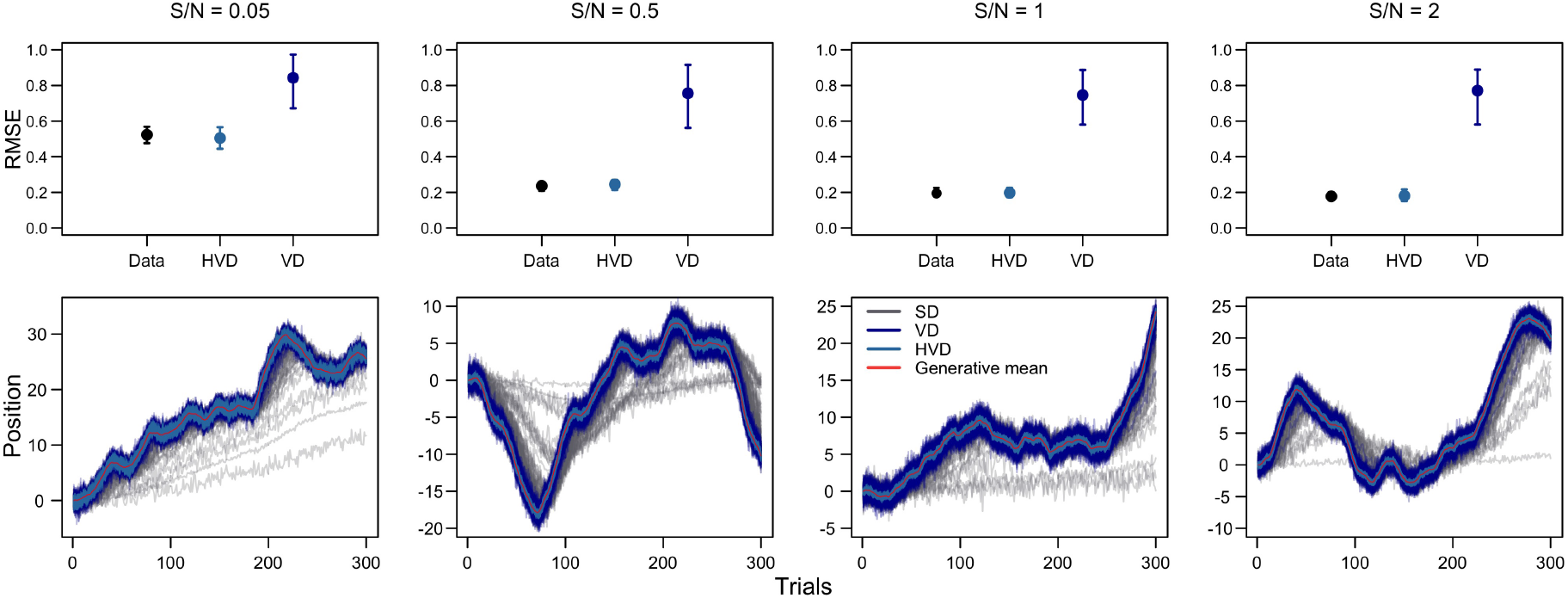
Predictions for a new subject on each experimental condition. Top panels represent the predicted RMSE for VD and HVD models compared to the actual RMSE of participants. Error bars correspond to the interquartile range and dots to the medians. RMSE for the SD model was excluded as it considerably increased the scale of the plot. Bottom panels correspond to trial-by-trial predictions of SD, VD, and HVD models for a new sequence of observations.

In summary, we observe from figures 6, 7 and 8, that incorporating the estimation of velocity into the standard reinforcement learning model increases its descriptive adequacy in a meaningful way. By additionally assuming a hierarchical structure we are able to generalize to new data sets and potentially to untested values of S/N while still providing an accurate description of data.

## Model comparison

Although RMSE provides a useful measurement of how accurately models can recover actual data, it cannot be used as a metric of model comparison as it ignores complexity, and generally more complex models will capture data better. In order to overcome this limitation, we implemented two standard methods in Bayesian modeling that incorporate model complexity into their computation. The first, is leave-one-out cross-validation (LOO), and the second, the Widely Applicable Information Criterion (WAIC). These techniques compute a pointwise estimator of the predictive accuracy of models for all data points taking one at a time. On the one hand, LOO is a type of cross-validation where the training dataset (the one used to tune the model) consist of all observations but one, which forms the validation dataset (the one to be predicted). In particular, we used a new approach to LOO developed by Vehtari et al. (2017) where a Pareto distribution is implemented to smooth weights in an importance sampling procedure, and that the authors termed PSIS-LOO (for Pareto smoothed importance sampling). On the other hand, WAIC works as an estimate of out-of-sample deviance which overcomes previous limitations of the deviance information criterion (DIC). Unlike DIC, WAIC is based on the entire posterior distribution and is valid under non Gaussian assumptions. In practice, PSIS-LOO and WAIC can be easily computed using the R package loo (Vehtari et al., 2017) and the log-likelihood evaluated at the posterior simulations of the parameter values (for more details on the computation of PSIS-LOO and WAIC, see equations 3, 10 and 11 of Vehtari et al. (2017)). These two methods return a measurement of deviance at predicting a new dataset and penalize model complexity. Differences of PSIS-LOO and WAIC between each model and the model with the lowest values for each metric are reported in table 2. More positive values indicate worse predictive performance. It is clear that HVD outperforms the other models according to both metrics. As expected, the SD has the worst predictive performance of the three models. It is important to note that model complexity in PSIS-LOO and WAIC is not defined in terms of parameter counts like in AIC or BIC. Instead, these metrics consider the variability of posterior predictions. When models make a wide range of predictions they are automatically penalized for the poor ones as in Bayes Factors. This is a relevant feature as in hierarchical models like HVD usually increasing the number of parameters reduces the variability of the predictions and, therefore, the complexity of the model (Lee & Vanpaemel, 2018).

**Table 2.** Differences of PSIS-LOO (Δ_PSIS–LOO_) and WAIC (Δ_WAIC_) between each model and the model with the lowest value for each metric. Higher values indicate worse predictive performance.

| Model | $\Delta_{PSIS-LOO}$ | $\Delta_{WAIC}$ |
| --- | --- | --- |
| SD | 92740 | 92467 |
| VD | 186 | 184 |
| HVD | **0** | **0** |

## Discussion

Humans and other animals often face environments that change over time. In some situations, these changes may occur gradually following a rate, and, in order to make accurate predictions, individuals should have a good estimate of this variable. However, the rate of change may not be readily inferred when peoples’ observations are corrupted with random fluctuations. In this paper, we tested people’s predictions in an environment with these characteristics by using a perceptual-decision making task. In our experiment, subjects predicted the future location of a spaceship that moved at a variable velocity and was corrupted with different levels of Gaussian noise. Our results show participants were able to predict the most likely future location of the spaceship with accuracy increasing for less noisy conditions. A standard reinforcement learning model (SD) was unable to qualitatively describe these results, and Bayesian inference showed learning rates reach values close to one in all conditions. This strategy is optimal only for a deterministic task and useful after the environment suffers abrupt and unpredictable changes, but inaccurate in a probabilistic setting that is changing gradually over trials. In an attempt to capture deviations from participants’ predictions, SD model assumes decision noise is high. This is likely to happen when a model is ignoring a crucial signal in data (namely the velocity component) and construing it as random variations. By incorporating a velocity term to the standard delta-rule (VD model), we were able to describe data in a more reasonable fashion. Furthermore, Bayesian inference showed that learning rates for velocity and decision noise increase and decrease, respectively, with the S/N. The above suggests that, in general, subjects updated their estimate of velocity faster and had less noisy predictions when observations were less corrupted by noise.

A hierarchical extension of VD suggests that the overall behavior of learning rates for velocity and decision noise can be modeled using a hyperbolic function. This model was able to capture participants’ errors as accurately as its non-hierarchical counterpart and to make reasonable predictions about a new participant. A formal model comparison using PSIS-LOO and WAIC also shows that the hierarchical model has the best predictive performance. Although not shown, this model additionally allows to make predictions about the overall behavior of parameters for untested values of S/N. This is because two of the model parameters (learning rates for position and memory retention for velocity) were invariant to our S/N conditions and therefore their values for other S/N conditions may be assumed to come from the same experiment-distributions. On the other hand, the hyperbolic function inferred for the learning rates of velocity and decision noise can take practically any positive value of S/N as input and provide an overall prediction of parameter values.

This work accords with other studies that propose humans are sensitive to higher-order variables that control the dynamics of the environment (Behrens et al., 2007; Courville, Daw, & Touretzky, 2006; McGuire, Nassar, Gold, & Kable, 2014; Meder et al., 2017; Ricci & Gallistel, 2017; Wittmann et al., 2016; Yu & Dayan, 2005). In particular, our model suggests that when the environment is changing smoothly at a variable velocity, subjects have an estimate of this quantity and use it to make predictions. Furthermore, we showed that this process is influenced by the expected uncertainty in the environment (Bland et al., 2012; Yu & Dayan, 2005), which enables faster learning of velocity when observations are less noisy (O’Reilly, 2013). Additionally, incorporating the velocity term allows to overcome the previous limitation of making the learning rate high when trial-to-trial changes are large, therefore reducing the influence of unpredictable variations caused by noise. The above is evident in figure 6 as the values of the learning rate for position and the decision noise are reduced by incorporating the estimate of velocity in the model, which, in turn, provides a much better description of data.

Although reinforcement learning models are common in tasks of belief updating in changing environments (Behrens et al., 2007; Nassar et al., 2010; Speekenbrink & Shanks, 2010; Wilson et al., 2013), some studies suggest that this process may not take place on a trial-by-trial basis as suggested by delta-rule models either when the environment suffers abrupt (Gallistel et al., 2014; Robinson, 1964) or gradual changes (Ricci & Gallistel, 2017). Instead, these works suggest that people follow a step-like pattern, some times updating their estimates after hundreds of trials have passed. This is true when people infer the parameter of a Bernoulli distribution (Gallistel et al., 2014; Ricci & Gallistel, 2017), however, the conditions under which people follow this pattern or a trial-by-trial update are not clear yet. Of particular interest to our paper is a recent error-driven approach to adaptive behavior using a control theoretic model known as PID (proportional-integral-derivative controller, Ritz et al. 2018). This model incorporates to the standard delta rule (proportional part), a weighted sum of the history of errors (integral part) and the difference between the current and previous error (derivative part). It is worth noting that the PI part of the model is algebraically equivalent to our VD model. A simple rearrangement of the integral part as a Markov process provides the update equation of the velocity term in the VD model. We believe our approach is computationally less expensive as it does not require estimating the full history of errors and their corresponding weights on every trial, but only the previous estimate of change and the current error. However, it is important to note that the model proposed here would perform poorly in task with abrupt changes of position or velocity as it suffers from the same pitfalls of models with fixed learning rates. PID model ameliorates this concern by incorporating the derivative part, which allows for sudden corrections when the model estimates depart from the generative process.

In summary, in this work we have provided evidence that people can use prediction errors to update an estimate of the rate of change when the environment is varying gradually over trials, and to update this quantity faster when observations are more reliable. Additionally, we have shown that a hierarchical Bayesian approach provides benefits in terms of predictive power and generalization. Finally, our results are in line with evidence that people and other animals can learn about higher order statistics of their environments and use that information to guide predictions.

## Acknowledgments

Supplementary material of this article, including code and data, is available as a project page on the Open Science Framework at https://osf.io/d6tjw/. This research was supported by the project PAPIIT IG120818. A preliminary version of this work was presented at the 51^*st*^ Annual Meeting of the Society for Mathematical Psychology in 2018.

